# Deaza-modification of MR1 ligands modulates recognition by MR1-restricted T cells

**DOI:** 10.1101/2022.05.11.491531

**Authors:** Haihong Jin, Nicole A. Ladd, Andrew M. Peev, Gwendolyn M. Swarbrick, Meghan Cansler, Megan Null, Christopher T. Boughter, Curtis McMurtrey, Aaron Nilsen, Karen M. Dobos, William H. Hildebrand, Deborah A. Lewinsohn, Erin J. Adams, David M. Lewinsohn, Melanie J. Harriff

## Abstract

MR1-restricted T (MR1T) cells recognize microbial small molecule metabolites presented on the MHC Class I-like molecule MR1 and have been implicated in early effector responses to microbial infection. As a result, there is considerable interest in identifying chemical properties of metabolite ligands that permit recognition by MR1T cells, for consideration in therapeutic or vaccine applications. Here, we made chemical modifications to known MR1 ligands to evaluate the effect on MR1T cell activation. Specifically, we modified 6,7-dimethyl-8-D-ribityllumazine (DMRL) to generate 6,7-dimethyl-8-D-ribityldeazalumazine (DZ), and then further derivatized DZ to determine the requirements for retaining MR1 surface stabilization and agonistic properties. Interestingly, the IFN-γ response toward DZ varied widely across a panel of T cell receptor (TCR)-diverse MR1T cell clones; while one clone was agnostic toward the modification, most displayed either an enhancement or depletion of IFN-γ production when compared with its response to DMRL. To gain insight into a putative mechanism behind this phenomenon, we used *in silico* molecular docking techniques for DMRL and its derivatives and performed molecular dynamics simulations of the complexes. In assessing the dynamics of each ligand in the MR1 pocket, we found that DMRL and DZ exhibit differential dynamics of both the ribityl moiety and the aromatic backbone, which may contribute to ligand recognition. Together, our results support an emerging hypothesis for flexibility in MR1:ligand-MR1T TCR interactions and enable further exploration of the relationship between MR1:ligand structures and MR1T cell recognition for downstream applications targeting MR1T cells.

## Introduction

As one branch of the adaptive immune system, T cells are responsible for mounting an immune response against specific infections. In the classical immunological axis, human major histocompatibility complex (MHC) cell surface proteins in complex with β-_2-_microglobulin (β_2_m), denoted as HLA-I molecules, are responsible for presenting pathogen-derived peptidic antigens on the cell surface. These complexes are sampled by T cells bearing αβ T cell receptors (TCRs), and when a cognate MHC-TCR complex is formed, signal transduction leads to a functional response by the T cell. The combination of the MHC allele, antigen, and TCR sequence therefore determine T cell reactivity. In contrast with this canonical T cell-mediated immune axis, HLA-Ib MR1-restricted T (MR1T) cells recognize small molecule metabolite antigens bound to MHC-I-related protein 1 (MR1) in complex with β_2_m. MR1 is a monomorphic protein bearing the same fold as classical MHC, though the antigen binding cleft displays unique biophysical properties. The groove is an “aromatic cradle” suited to binding small heterocyclic compounds including secondary metabolites generated during riboflavin biosynthesis [1, 2] a pathway that is present in some bacterial and fungal species but not higher order eukaryotes. Cell surface presentation of agonist metabolites by MR1 therefore signals the presence of microbes to cognate MR1T cells, which occupy a unique niche in the immune recognition of microbial infection.

A subset of MR1T cells known as mucosal-associated invariant T (MAIT) cells are prevalent in human mucosal tissues and blood [3] and have been implicated in early immune responses to numerous microbial infections in both mice and humans [4]. In humans, MAIT cells express TCRs that employ limited diversity in the β chain and are restricted to usage of the TRAV1-2 gene rearranged with a small number of TRAJ genes (TRAJ20 and TRAJ33). This encodes a TCR with a conserved tyrosine residue (typically Y95) in the CDR3α loop which is responsible for forming a critical hydrogen bond with the 2’-hydroxyl of the ribityl moiety of the MR1-bound antigen [5-8]. Mutation of Y95 disrupts the ability of MAIT cell clones to respond to stimulatory ligands and Y95F has been shown to decrease the affinity of MR1:ligand-TCR interactions by an order of magnitude [7, 8]. Some ligands also promote contacts with the CDR3β loop of the TCR, though the significance of these interactions, and those between CDR3β and MR1 itself, are unclear [7, 9]. More recently, MR1T cells utilizing alternate TRAV genes have been described [10, 11], some of which have significantly different footprints on the MR1 groove and, by nature of alternative TRAV genes, different ligand recognition mechanisms than typical MAIT TCRs [9, 10]. Further, evidence continues to build that TCR-diverse MR1T cells can distinguish between different ligands [10-14], suggesting that differences in ligand structures could result in the expansion of distinct subsets of MR1T cells. While the monomorphic nature of MR1 makes MR1T cells a tempting target for adoptive T cell therapies [15], this increasing evidence for MR1T cell diversity makes understanding strategies for molecular recognition of antigens a prerequisite to harnessing their therapeutic potential.

MR1T cell agonists are generally found in two classes of riboflavin metabolites, the pyrimidines and the ribityllumazines, though additional ligands continue to be identified [13, 14, 16]. Pyrimidines, such as 5-(2-oxopropylidenamino)-6-D-ribitylaminouracil (5-OP-RU), the most potent MR1T cell-activating ligand identified to date, are secondary metabolites formed from the spontaneous interaction between the riboflavin precursor 5-amino-6-D-ribitylaminouracil (5-A-RU) and other microbial- or mammalian cell-derived metabolites such as methylglyoxal (in the case of 5-OP-RU) [1]. The pyrimidines are unique in that they are able to form a Schiff base with the K43 residue of MR1 [1], which is buried deep in the ligand binding pocket, though the bond is not required for MR1T cell activation. These pyrimidines, however, are intrinsically unstable in solution because they undergo spontaneous ketone/amine condensations under physiological conditions [1, 17]. This reaction results in the production of ribityllumazines, which are bicyclic, aromatic structures with a conserved lumazine base but variable adducts at C6 and C7 depending on the metabolite with which 5-A-RU condenses [1]. One such ribityllumazine, 6,7-dimethyl-8-D-ribityllumazine (DMRL), can also be formed in a biosynthetic cycle wherein microbial lumazine synthase (part of the riboflavin biosynthesis pathway) condenses 5-A-RU and 3,4-dihydroxy-2-butanone-4-phosphate [1] through a putative pyrimidine intermediate, 5-(1-methyl-2-oxopropylideneamino)-6-D-ribitylaminouracil (5-MOP-RU) (Figure 1). Known ribityllumazines are not expected to form a Schiff base with the K43 residue of MR1. These ligands are typically less potent MR1T cell agonists than the pyrimidines [9], but the molecular basis of MR1T cell antigen potency and the role of different classes of ligands *in vivo* is unclear. It is obvious, however, that 5-A-RU is a critical component for MR1 ligand generation in many microbes [1, 13, 18, 19] and that determining the contribution of ribityllumazines to MR1T cell biology is a critical step toward leveraging this immune axis for immunotherapeutic applications.

**Figure 1.**
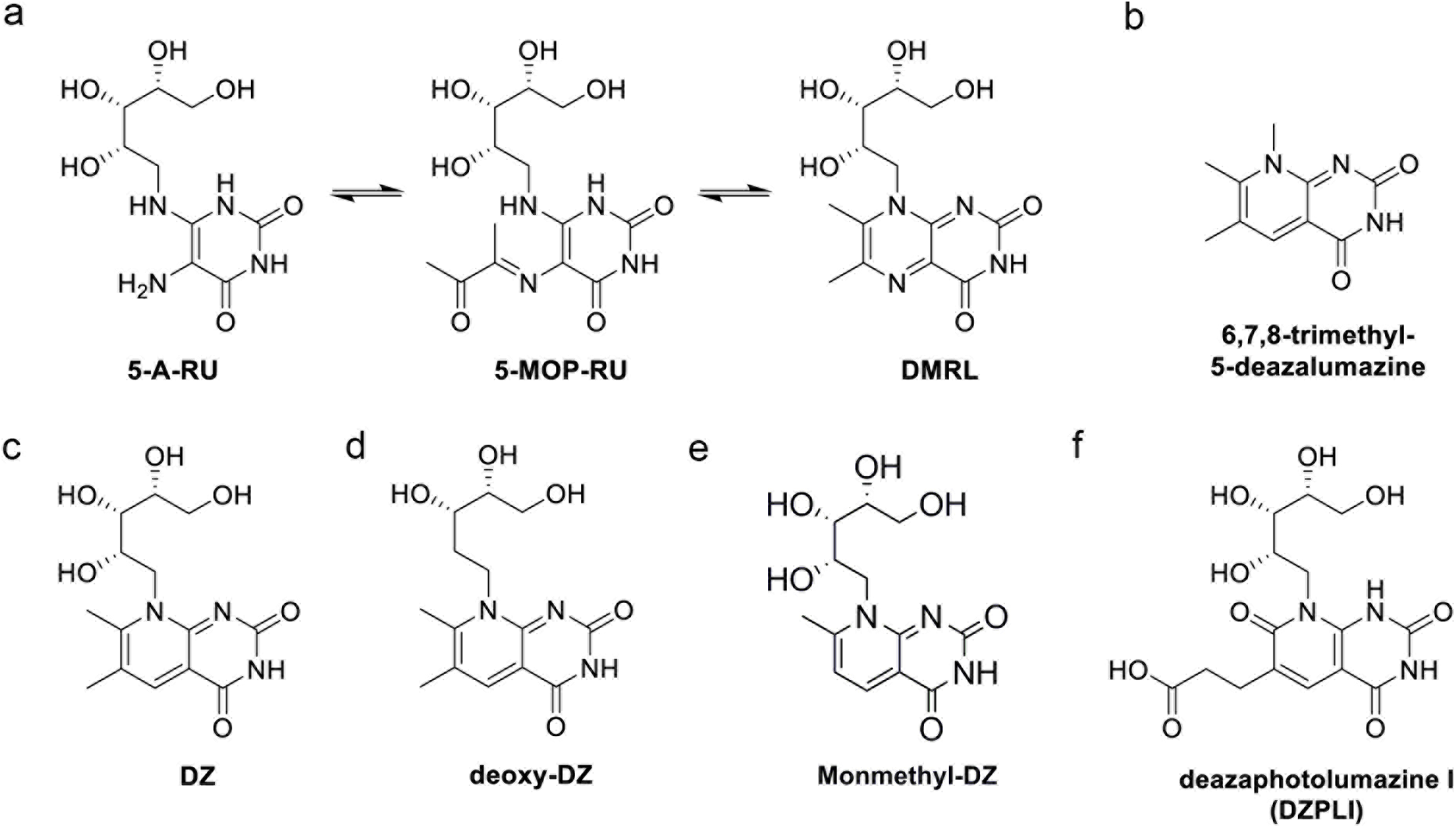
Proposed chemical structures for deaza-molecules. a) Reaction scheme from 5-A-RU to DMRL indicating reversible reactions via ketimine hydrolysis. b) Non-ribityl portion of the deaza-version of the lumazine intermediate. c) Proposed DZ chemical structure. d) Proposed 2’-deoxy-DZ chemical structure. e) Proposed monomethyl-DZ chemical structure. f) Proposed DZ-PL1 chemical structure.

The abundance of MR1T cells in tissues, and the ability of MR1T cells to be stimulated by a novel and conserved class of molecules produced by a broad range of pathogens, warrants a better understanding of the relationship between ligand structure and MR1T cell activity. Here, we sought to assess MR1T cell ligand selectivity using modified deaza-analogues of known ribityllumazine MR1T cell agonists. We assessed the ability of these analogues to activate MR1T cells and stabilize MR1 on the cell surface, then used *in silico* techniques to model how minor ligand modifications may contribute to our cellular observations. We found that the deaza-analogue of DMRL was recognized differentially across a panel of TCR-diverse MR1 cell clones. In silico molecular docking and molecular dynamic simulations further demonstrated surprising differences possibly contributing to this differential recognition. Together our results support the continued study of interactions between MR1T cell ligands in the context of MR1:ligand-TCR complexes.

## Results

### Synthesis of deazalumazine (DZ), 2’-deoxy-deazalumazine (2’-deoxy-DZ), monomethyl deazalumazine (monomethyl-DZ) and deazaphotolumazine I (DZPLI)

In order to synthesize deaza-forms of lumazines, we took inspiration from preparations of lumazine synthase inhibitors where the 5-nitrogen of the non-ribityl lumazine core was replaced with a carbon [20] (Figure 1a). Specifically, we replaced of the 5-nitrogen of 6,7-dimethyl-8-D-ribityllumazine (DMRL) and 7-methyl-8-D-ribityllumazine with a carbon to produce the known compounds 6,7-dimethyl-8-D-ribityldeazalumazine (deazalumazine or DZ) and 7-methyl-8-D-ribityldeazalumazine (monomethyl-DZ), respectively (Figure 1b and 1c). Additionally, we prepared 2’-deoxy-DZ (Figure 1d), lacking the 2’-hydroxyl of the ribityl moiety, and a deaza-version of photolumazine I (DZPLI, Figure 1e), a novel MR1T cell-activating ribityllumazine ligand we recently described [13]. It is worth noting that the original design was to create a 5-deaza version of 5-OP-RU, but this compound spontaneously cyclized during synthesis to produce monomethyl-DZ.

Syntheses of analogues DZ, 2’-deoxy-DZ, DZPLI and monomethyl-DZ are summarized in Figure 2. Substitution of commercially available 6-chlorouracil **1** with ribitylamine or 2’-deoxyribitylamine in water at 150°C in a microwave reactor led to the corresponding ribityluracil **2a** or 2’-deoxyribityluracil **2b** (Figure 2a). The use of a microwave reactor rather than conventional heating boosted the reaction yield 10-fold. The ribityluracil compounds **2a** and **2b** were then reacted with the sodium salt of 2-methyl-3-oxo-butanol under acidic conditions at 100°C to give DZ and 2’-deoxy-DZ, respectively. DZPLI was formed by reacting ribityluracil **2a** with 1,5-diethyl-2-formylpentanedioate under acidic conditions at 100°C. Monomethyl-DZ was prepared in four synthetic steps as illustrated in Figure 2b. The initial step, formation of aldehyde **4**, was accomplished via formylation of commercially available 4-chloro-2,6-dimethoxypyrimidine **3** by direct ortho-lithiation with n-butyllithium in tetrahydrofuran (THF) followed by formylation with N,N-dimethylformamide (DMF). Subsequently, the Wittig reaction was accomplished by reacting aldehyde **4** with commercially available (acetylmethylene) triphenylphosphorane; the product **5** was obtained in high yield. Compound **6** was obtained by chloro-displacement of compound **4** with ribitylamine. The crucial part of the synthesis was the acidic removal of the compound **6** methoxy groups to give monomethyl-DZ. After screening a variety of conditions, e.g., BBr_3_ in CH_2_Cl_2_ and various concentrations of HCl, it was found that 6 M hydrochloric acid was able to convert the pyrimidine ring to the corresponding uracil with acceptable yield. Using HPLC analysis, the purities of DZ, 2’-deoxy-DZ, monomethyl-DZ and DZPLI were determined to be greater than 95%.

**Figure 2.**
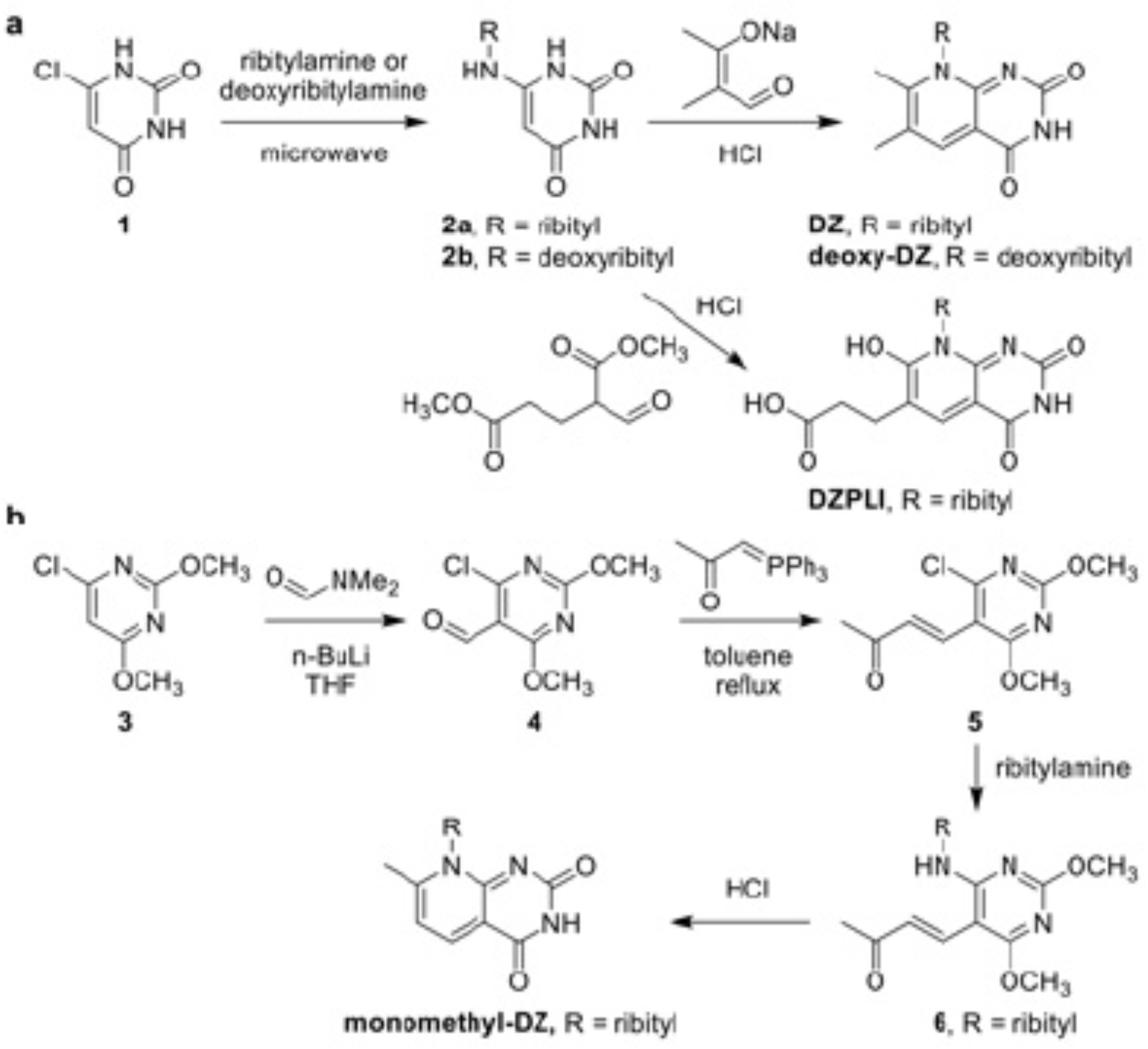
Synthesis of deazalumazine (DZ), 2’-deoxy-deazalumazine (2’-deoxy-DZ), and deazaphotolumazine I (DZPLI). a) Chemical synthesis scheme for DZ, 2’-deoxy-DZ, and DZ-PLI. b) Chemical synthesis scheme for monomethyl-DZ. Synthesis details for both schemes are described in the Methods.

### MR1T cell responses to DZ

To determine whether or not DZ is a MR1T cell antigen, we tested the synthetic DZ molecule, alongside DMRL, a known MR1 agonist ligand and the DZ parent structure, for the ability to stimulate a TCR-diverse panel of MR1T cell clones in an IFN-γ ELISPOT assay with human monocyte-derived dendritic cells (DC) as the antigen presenting cell. D426G11, D481A9, D481C7, and D481F12 are previously described TRAV1-2^+^ MR1T (MAIT) cell clones where the TRAV1-2 α-chain is paired with diverse β-chains and CDR3 amino acid sequences (Table 1, Figure 3a) [12, 13]. We also tested a non-traditional MR1T cell clone, D462E4, which expresses TRAV12-2 paired with a unique β-chain (Figure 3a)[11]. Despite the minor 5N to 5C substitution, the responses of the MR1T cell clones to DC pulsed with DZ varied widely between clones, and compared to the response to DMRL (Figure 3a). There were no significant differences in the responses to DMRL or DZ for D481F12 (p=0.91). D426G11 responded to DZ, but significantly less well than to DMRL (p<0.0001). The response of D481A9 to DMRL was also significantly higher than to DZ (p<0.001); in fact, this clone did not respond to DZ at any of the tested concentrations. In contrast, D481C7 had a significantly more robust response to DZ than to DMRL (p=0.0007). Interestingly, D462E4, which does not recognize DMRL at these concentrations, responded significantly more robustly to DZ (p=0.0008). These data suggest that DZ is capable of acting as an MR1T cell agonist, and that a minor chemical substitution that is not expected to be TCR-accessible is nonetheless able to modulate MR1T cell responses.

**Figure 3.**
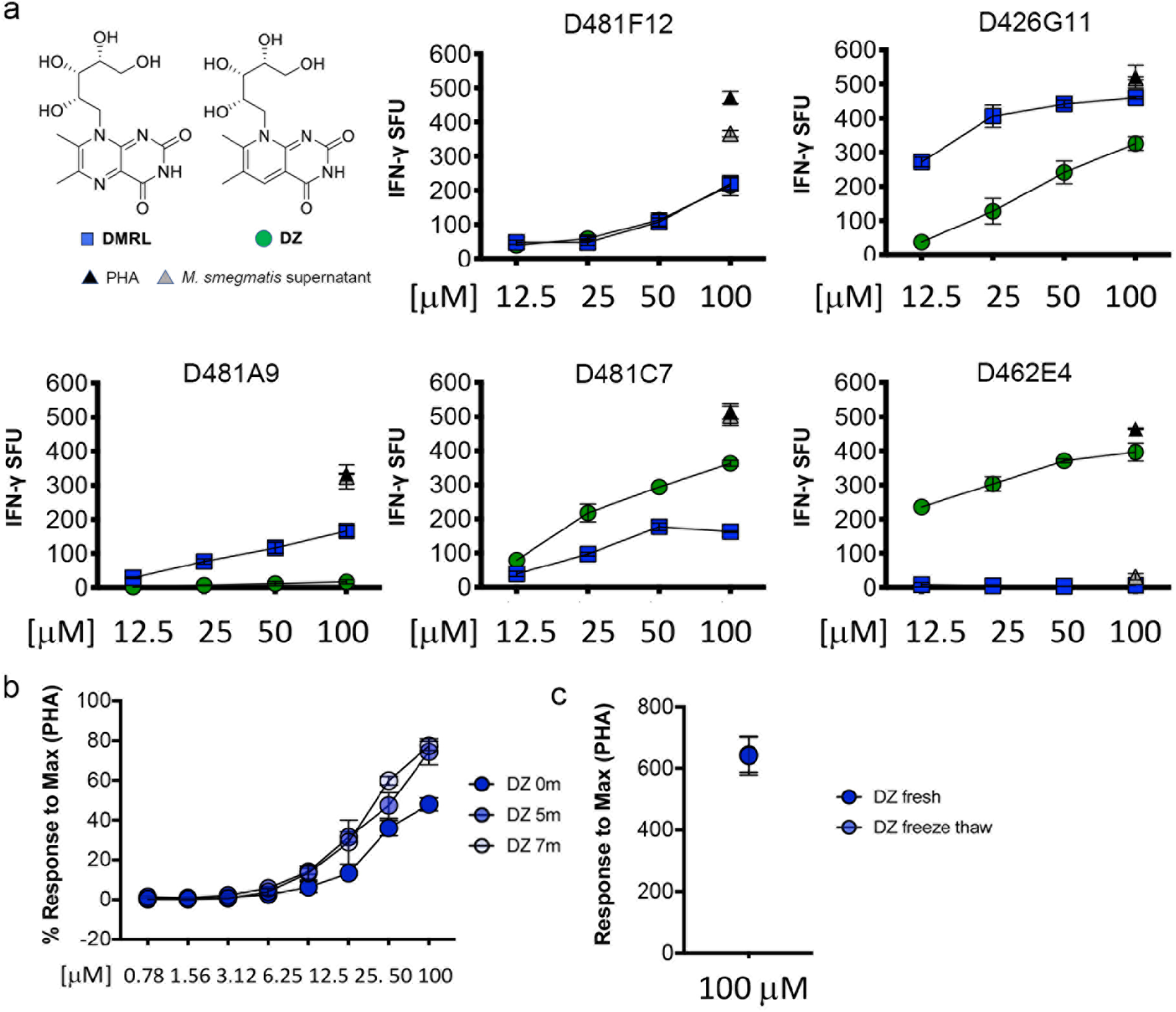
MR1T cell responses to DZ. a) The IFN-γ response by MR1T cell clones was measured in an ELISPOT assay with 1×10^4^ DC pulsed with the indicated concentrations of DMRL or DZ. P values were determined by simple linear regression of the slope and elevation of for DZ and DMRL. b) The same synthesis of DZ was tested as described in a) following storage of the compound in dimethylsulfoxide (DMSO) at -80°C for 5 and 7 months, using D481 C7 response as the readout. c) A single vial of DZ stored in DMSO at -80°C was tested initially after thawing or following a cycle of refreezing and thawing prior to use in an ELISPOT assay, using D481 C7 response as the readout. Phytohemagglutinin (PHA) and supernatant from *M. smegmatis*, as described in the Methods, were used as positive controls for all assays. Error bars indicate the standard deviation of technical replicates. Data shown are representative of 3 independent experiments.

To confirm the expected functional stability of DZ, we measured MR1T cell responses to DC pulsed with different synthesis batches of DZ at multiple time points after synthesis, and also after freeze-thawing the DZ. Two separate synthesis batches of DZ were tested upon delivery and 5-7 months after suspension in DMSO and storage at -80°C. For both batches, there was no loss of MR1T cell activity in response to freshly synthesized DZ versus that which had been in the freezer (Figure 3b). Additionally, there was no loss of activity following an additional freeze-thaw cycle prior to the ELISPOT assay (Figure 3c). Together, these results confirm DZ is stable when stored at -80°C and is resistant to degradation from freeze-thaw cycles.

### Impact of chemical modifications to DZ and PL1 on MR1T cell responses

2’-deoxy-DZ, monomethyl-DZ and DZPLI were then tested in an ELISPOT assay measuring IFN-γ production by the D481C7 MR1T cell clone, which was robustly stimulated by DZ and PLI. Removing the 2’-hydroxyl group from the ribityl chain of DZ to generate 2’-deoxy-DZ reduced the response of the D481C7 clone 10-fold (Figure 4a). This is consistent with other reports that have determined the importance of the 2’-hydroxyl group for MR1T TCR recognition [1, 5, 6, 8, 9]. In contrast, modifying DZ to generate monomethyl-DZ had almost no impact on the response of the D481C7 clone (Figure 4a). Previously, we demonstrated that only the D481C7 clone had robust responses to PLI, while the other clones had little to no response [13]. MR1T cell clones that did not recognize PLI also did not recognize DZPLI (data not shown). Whereas the D481C7 clone had a significantly higher response to the deaza-version of DMRL (DZ) when compared to DMRL itself, the response of the D481C7 clone to DZPLI was reduced nearly 5-fold when compared to PLI (Figure 4b). Together, these results show that small changes in the structure of MR1T cell ligands can dramatically alter the ability of these ligands to be recognized by MR1T cells in distinct ways, and support the hypothesis that MR1T TCRs can recognize very specific patterns.

**Figure 4.**
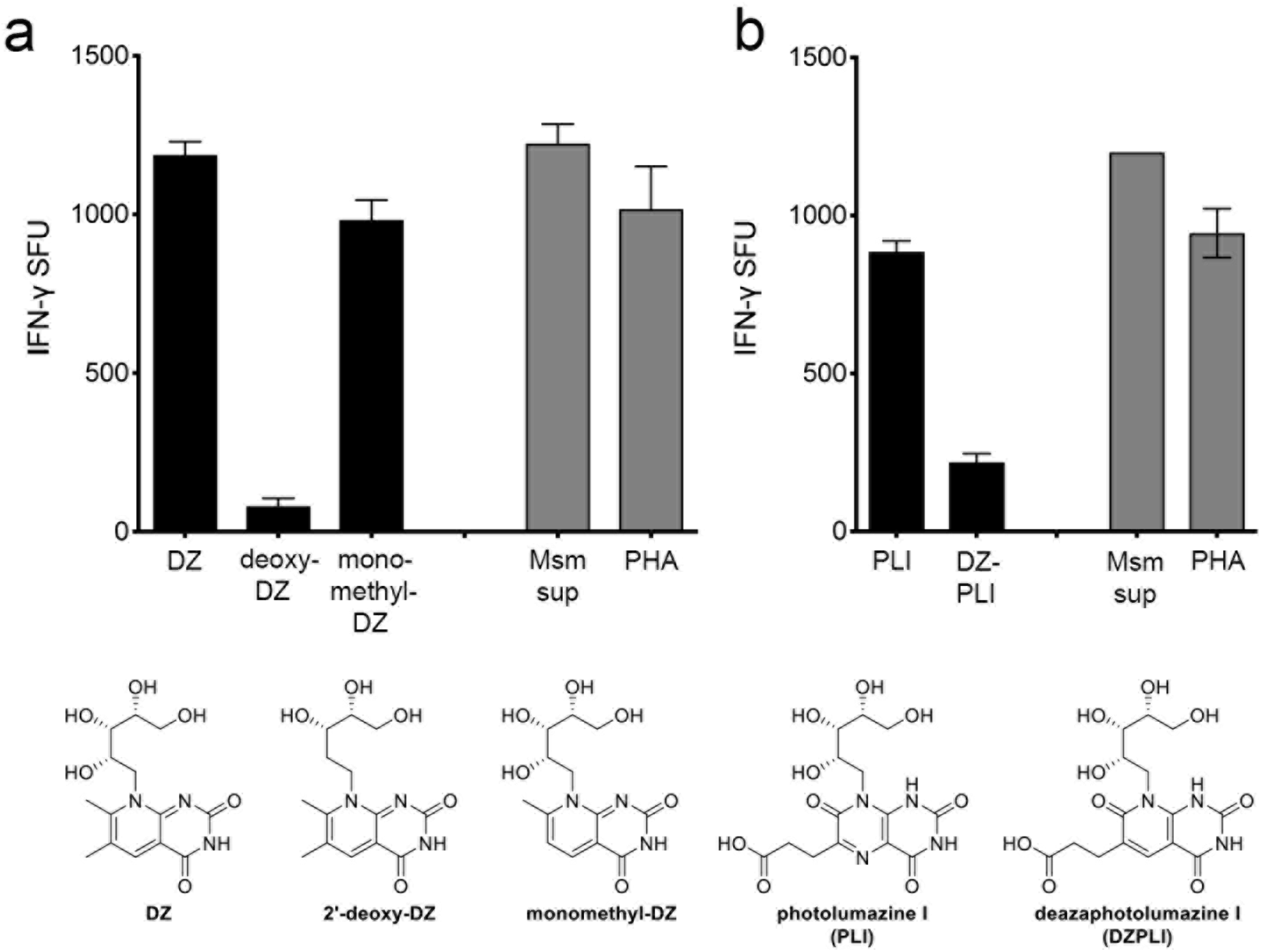
Impact of chemical modifications to DZ and PLI on MR1T cell responses. The IFN-γ response by D481C7 MR1T cell clones was measured in an ELISPOT assay with either 1×10^4^ (a) or 2×10^4^ (b) DC pulsed with a) DZ, 2’- deoxy-DZ, or monomethyl-DZ at 100 μM, and b) PLI or DZ-PLI at 50 μM as antigen presenting cells. PHA and supernatant from *M. smegmatis* were used as described as positive controls for all assays. Error bars indicate the standard deviation of technical replicates. Data shown are representative of 3 independent experiments.

### MR1T cell recognition of DZ is MR1-dependent

We next confirmed the requirement for MR1 in MR1T cell responses to DZ in two ways. First, we tested MR1T cell responses within the context of blocking with the α-MR1 26.5 blocking antibody. Here, when DC were incubated with the α-MR1 26.5 antibody prior to being pulsed with DZ, the response by both the D481C7 and D426G11 MR1T cell clones was completely abrogated (Figure 5a-b). Second, we tested MR1T cell responses in an MR1 knockout cell line. We first demonstrated that both MR1T cell clones could recognize DZ presented by BEAS-2B wild-type cells (Figure 5c). As expected based on previous experiments, the response of the D426G11 clone was less than that of the D481C7 clone to both DC and BEAS-2B cells. Although the response to DZ-pulsed BEAS-2B cells, a non-professional antigen presenting cell, was lower for both clones, MR1T cells are nonetheless clearly capable of recognizing DZ presented by these cells. When either MR1T cell clone was incubated with DZ-pulsed BEAS-2B:ΔMR1 cells, however, they were unable to produce IFN-γ (Figure 5d). Confirming the requirement for MR1, the IFN-γ response by both MR1T cell clones was restored when MR1 expression was reconstituted in the BEAS-2B:ΔMR1 cells (Figure 5d). Together, these results demonstrate that, as with other ligands including DMRL, professional antigen presenting cells and airway epithelial cells are capable of presenting DZ to MR1T cells in an MR1-dependent manner. These data demonstrate that DZ is a bona fide MR1T cell antigen and contribute to the realization of ligand discrimination among distinct MR1T TCRs.

**Figure 5.**
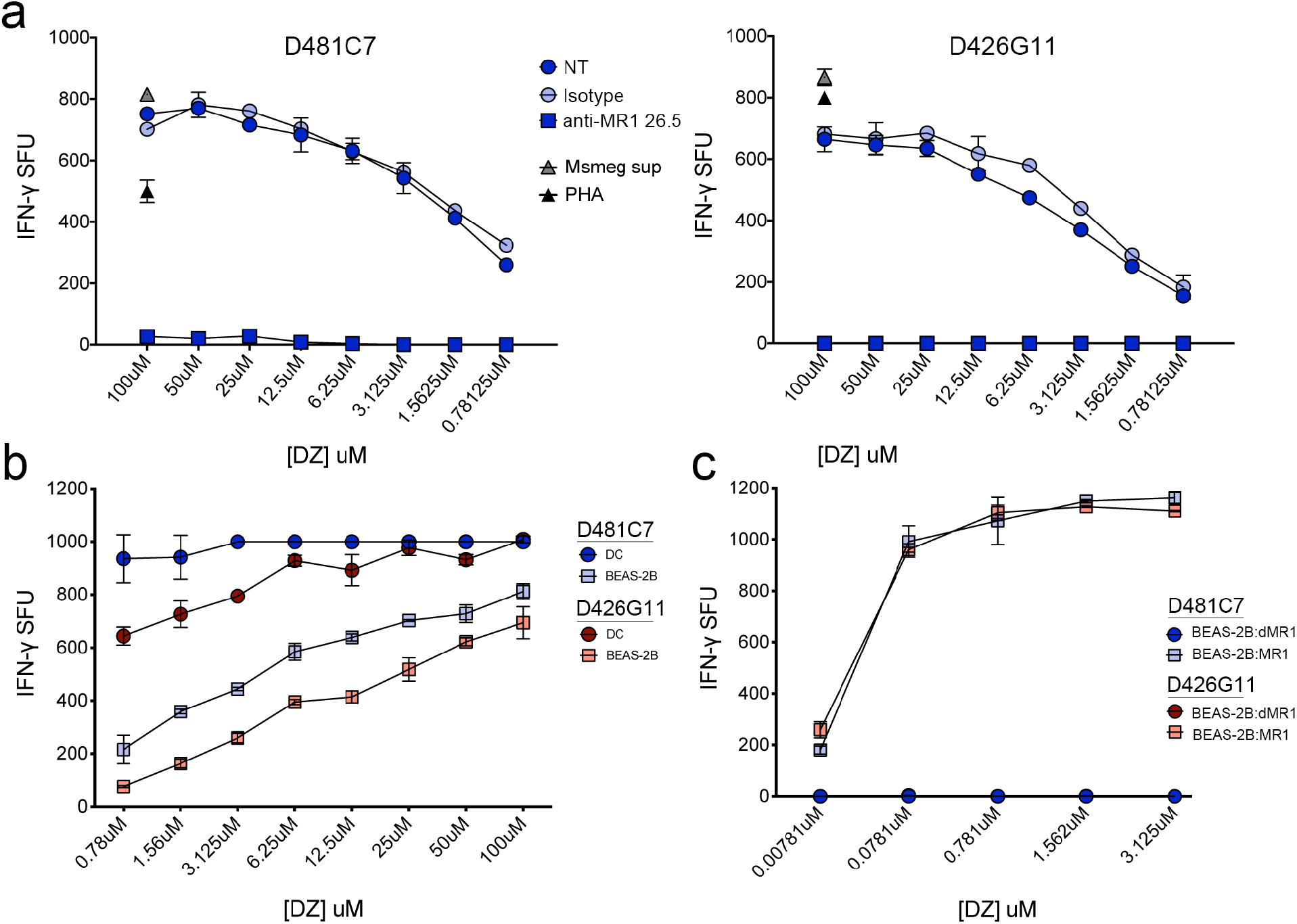
MR1T cell recognition of DZ is MR1-dependent. A-b) The IFN-γ response by D481C7 (a) or D426G11 (b) was measured in an ELISPOT assay with 1×10^4^ DC pulsed with DZ at the indicated concentration. Where indicated, the α- MR1 26.5 antibody or isotype control was added one hour prior to adding DZ to the wells. PHA and supernatant from *M. smegmatis* were used as described as positive controls for the assay. c) The IFN-γ response by D481C7 or D426G11 was measured in an ELISPOT assay with 1×10^4^ DC or 1×10^4^ wild-type BEAS-2B cells pulsed with DZ at the indicated concentration. d) The IFN-γ response by D481C7 or D426G11 was measured in an ELISPOT assay with 1×10^4^ BEAS- 2B:ΔMR1 or BEAS-2B:ΔMR1+MR1 cells pulsed with DZ at the indicated concentration. Error bars indicate the standard deviation of technical replicates. Data shown are representative of 3 independent experiments.

### Ligand-dependent MR1 stabilization on the cell surface

In the absence of ligand, MR1 is sequestered in the ER and endosomal compartments, but ligand availability triggers egress to the cell surface through an unknown mechanism [21, 22]. Therefore, we looked at whether or not these synthetic ligands were capable of stabilizing MR1 on the cell surface, similar to what has been observed previously for the non-stimulatory ligand 6-formylpterin (6-FP) and 5-OP-RU [7, 22]. BEAS-2B cells overexpressing MR1 were incubated with saturating amounts of 6-FP, DZ, 2’-deoxy-DZ, monomethyl-DZ, PLI, or DZ-PL1 then surface stained for MR1 using the α-MR1 26.5 antibody. Here, we found that all of the ligands tested that were able to stimulate at least one MR1T cell clone (DZ, monomethyl-DZ, and PLI) were able to stabilize MR1 on the cell surface, whereas DZPLI, which minimally stimulated only the D481C7 MR1 clone, was unable to stabilize MR1 on the cell surface (Figure 6). Interestingly, 2’-deoxy-DZ, which also induced reduced MR1T cell activation, nonetheless stabilized MR1 on the cell surface (Figure 6) consistent with other findings [23].

**Figure 6.**
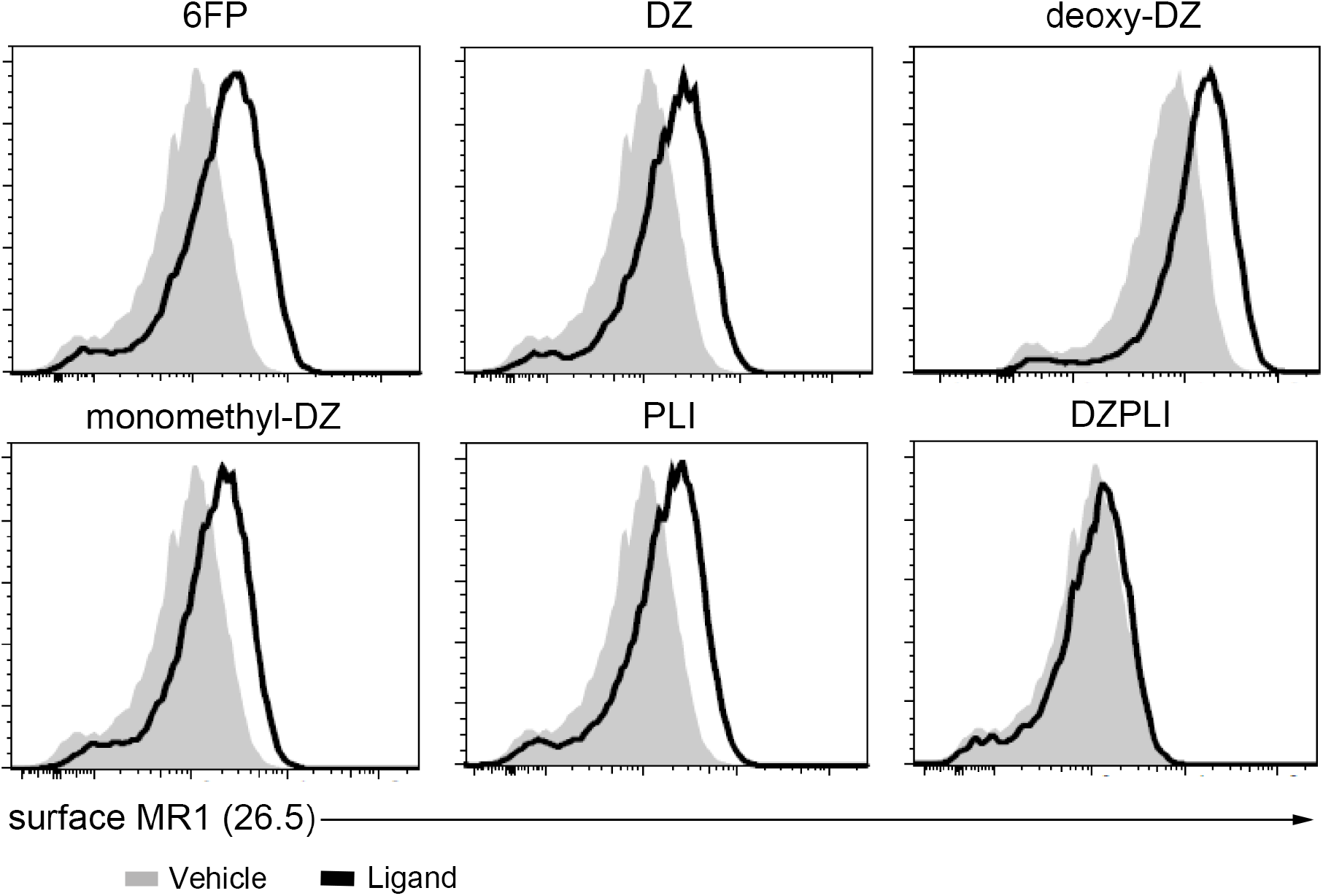
MR1T cell activating potential of MR1 ligands and MR1 stabilization on the cell surface. WT BEAS-2B cells overexpressing MR1-GFP were incubated with indicated ligands at 100μM for 16 hours. MR1 on the cell surface was detected by surface staining with the α-MR1 26.5 antibody conjugated to APC. Grey shaded histograms represent the vehicle and black solid histograms represent the ligand. Histograms are representative of 2 independent experiments.

### DMRL, DZ, and 2’-deoxy-DZ exhibit differential dynamics in the MR1 binding pocket

In order to gain insight into a possible mechanism behind the differential agonistic properties of these chemically-modified forms of DMRL, we employed structural and computational approaches to provide atomistic insights into our cellular observations. Limited ligand availability and the difficulty of refolding MR1 with ribityllumazines [24] made co-crystallization of the ligand with MR1 unfeasible, so AutoDock Vina [25] was used to model their docking. The original atomic coordinates of the β_2_m/MR1 complex were taken from the x-ray crystal structures (“donor structures”) of human MR1 and human β_2_m in complex with either the ligand 6-FP only (PDB ID: 4GUP [2]) or in complex with a MAIT TCR and the ligand 7-methyl-6-hydroxy-8-D-ribityllumazine (HMRL) (PDB ID: 4L4V [8]). β_2_m/MR1 coordinates were also isolated from 4LCC, a crystal structure of chimeric human-bovine MR1 in complex with rRL-6-CH_2_OH and a MAIT TCR; this structure was used to compare the crystallographically-determined and docked binding modes of a different ribityllumazine. Ligand coordinates for HMRL, DMRL, DZ, 2’-deoxy-DZ, monomethyl-DZ, PLI, DZPLI, and rRL-6-CH_2_OH were then drawn *de novo* using BIOVIA Discovery Studio [26]. We selected the A’ pocket of MR1 as the docking search space, as this is where the ribityllumazines and pyrimidines have been shown to bind the groove [1, 2, 5, 6, 8]. Vina then predicted the binding mode of each ligand based on calculated chemical potentials of all possible ligand conformations. Further details for this docking protocol can be found in the Methods section. Vina provides an internal scoring system to rank binding modes (“affinities” measured in kcal/mol), and the highest-ranking conformation was chosen for each ligand. Vina-scored affinities can be found in Table 2 and visualizations of these models are shown in Figure S1. Coordinates of the docks are provided in the supplementary material. While these Vina scores are given in kcal/mol, the scoring function of AutoDock Vina does not necessarily correlate well with experimental affinities, and instead should be considered as an internal reference score [25].

Docked ligands that have been previously crystallized show good agreement with experimentally determined structures, including docking HMRL back into 4L4V (Figure S2b). Notably, rRL-6-CH_2_OH docked back into 4LCC is flipped about the plane of the ring that is observed in the experimental crystal structure [5] (Fig S2a). However, docked rRL-6-CH_2_OH aligns with other ligand modes with respect to ring orientation and projection of the ribityl moiety (PDB IDs: 4L4V, 6PUF [8, 9]). Comparing the DMRL analogues in the 4GUP donor structure, the lumazine cores overlap generally, but DMRL is particularly aligned with DZ, presumably because the C6 and C7 adducts are identical for these two compounds (Figure S1a). More notably, the ribityl chain of 2’-deoxy-DZ (purple) is posed differently from DMRL, DZ, and monomethyl-DZ, likely owing to the absence of the 2’ hydroxyl group. Similar trends were seen for the DMRL analogues docked into 4L4V (Figure S1b). However, minor changes in ligand orientation were observed when comparing the dock of each ligand across the donor structures: the DMRL analogues docked into the 4L4V donor structure experience a slight rotation about C7, and ∼0.5 Å translation of each ligand, though they stayed in the same plane (Figure S1c; only DMRL is shown for clarity). This resulted in a shift into the pocket (with greater movement of the uracil ring) and reorganization of the ribityl moiety, which are likely due to the well-described structural adaptations of the MR1 binding groove to TCR binding (such as the rotation of MR1 residue Y152 and relaxation of the α_2_ helix, which together “pry open” the groove) [8]. We also attempted docking of PLI and DZPLI and observed that in the 4GUP donor structure, they were essentially superimposable (Figure S1d). The rings overlapped well with the DMRL analogues (not shown), but that on docking into 4L4V, they experienced differential rotation about the C7 and ring-bridging C-C bond, respectively (Figure S1e-f). These data suggest that this docking strategy is sensitive to minor changes in both protein and ligand structure, lending confidence toward these models.

To validate the docking mode of the DMRL analogues, we employed all-atom molecular dynamics of the MR1:ligand complexes to simulate the temporal evolution of each ligand in the binding pocket. We performed these simulations on the 4GUP docks to prevent bias of the starting model based toward protein alterations upon TCR engagement (as seen in 4L4V). Simulation details can be found in the Methods section. Starting from the structures of each ligand docked into the 4GUP donor structure, we simulated each system for 80 nanoseconds, running simulations in triplicate to minimize stochastic effects from standard velocity initialization [27, 28]. Simulations show that MR1:ligand complexes are stable (as measured by ligand RMSD) across time (Figure S3), adding further confidence to our docked models.

Given the similarity of the DZ and 2’-deoxy-DZ binding modes, we reasoned that the inability of 2’-deoxy-DZ to stimulate MR1T cell clones (Figure 4), while still being able to stabilize MR1 at the cell surface (Figure 6), was due to changes in TCR contacts with the ligand and/or MR1, as has been seen in crystallographic studies of deoxyribityl analogues of 5-OP-RU in complex with a MAIT TCR [9]. To visualize the proximity of DZ, and 2’-deoxy-DZ to MR1T TCR loops, we selected the frame 25 of a representative simulation for these ligands and aligned them to the crystal structures of ternary complexes with various ligand-TCR combinations (Figure 7e) (PDB IDs: 4IIQ, 4L4V, 4PJ7, 4PJ8, 4L9L [5-8]). This frame corresponds to 0.5 ns of equilibration and 0.25 ns of unrestrained all-atom dynamics and was chosen to highlight initial subtle changes in the docking before any potential larger shifts of the ligand location and conformation occur (Figure 7a-b). We were able to recapitulate the close contact of Y95 of each of the MAIT TCR CDR3α when aligning to DZ docked into 4GUP (data not shown), but after this equilibration, the 4’-hydroxyl of DZ was positioned more closely to CDR3α Y95 than was the 2’-hydroxyl (Figure 7c). In the 2’-deoxy-DZ alignment, none of the hydroxyl moieties were close enough to Y95 to make a competent polar contact (Figure 7d). Likewise, there were CDR3β sidechains from 3 TCRs within distance to make polar contacts with each of the docked ligands (data not shown), but after simulation, the conformation of the MR1:ligand complexes changed insomuch that only the 4PJ8 TCR was in proximity to do so. Since these are alignments with crystallographically-determined ternary complexes and not simulations of entire ternary complexes, the dynamics of MR1:ligand and TCR are not coupled. Thus, we wouldn’t expect this to directly reflect the “real” binding, but perhaps a close approximation.

**Figure 7.**
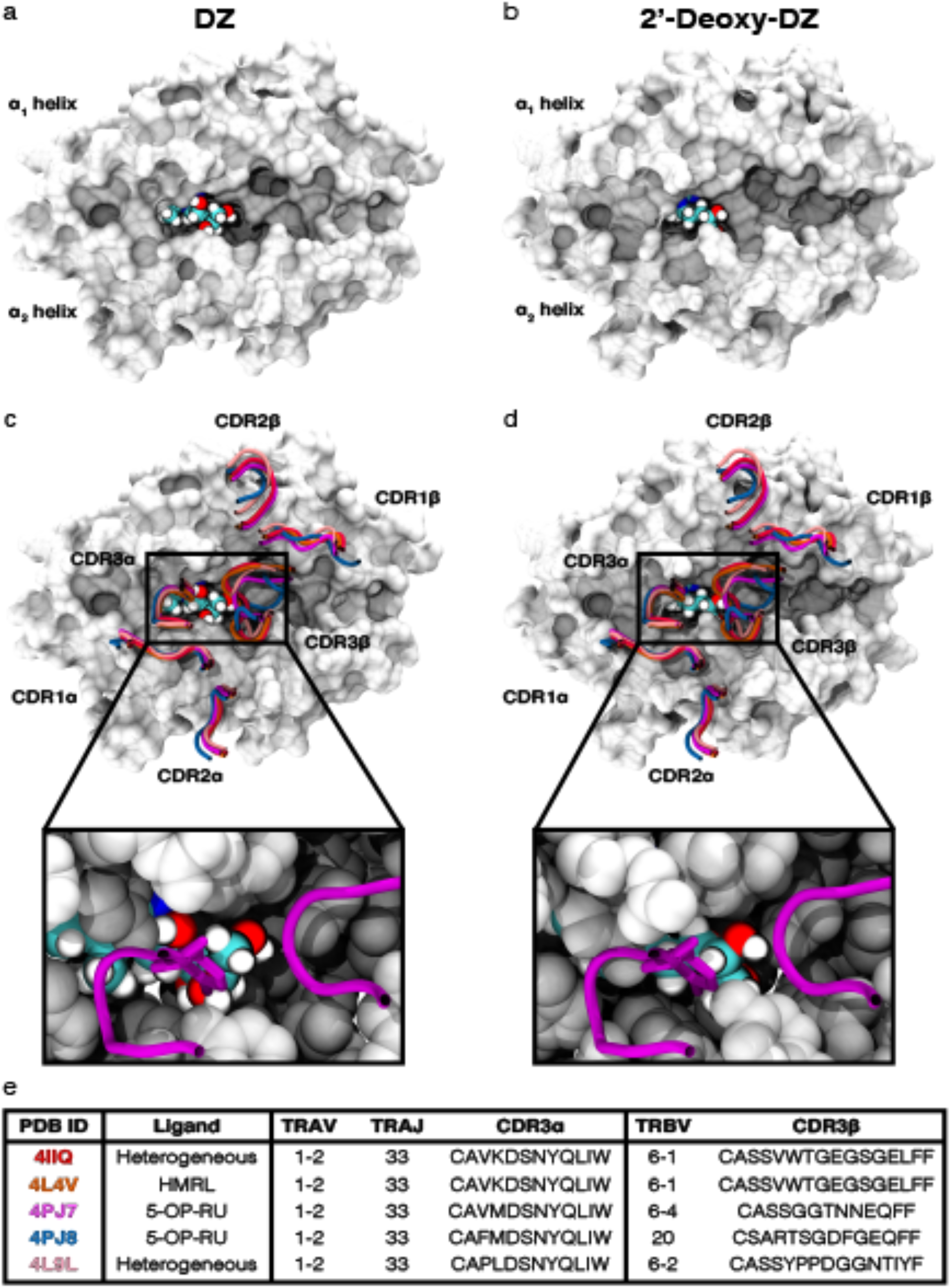
Visualizations of DZ and 2’-deoxy-DZ orientation in MR1, and their proximity to the TCR. a-b) DZ (a) and 2’- deoxy-DZ (b) ligand orientation at the 25^th^ frame of a representative simulation; the MR1 α1/α2 platform is represented as a surface and the ligand is represented by van der Waals-radius spheres. c-d) Overlay of MAIT TCR CDR loops with the selected frame from DZ (c) and 2’-deoxy-DZ (d). The inset is a zoomed-in representation of a TCR (PDB ID: 4PJ7) with the sidechain of Y95 shown explicitly. This is meant as a visual guide to aid in discussion of differential TCR interactions with MR1:ligand.

Interestingly, over the course of these simulations, we found that DMRL, DZ, and 2’-deoxy-DZ adopt differential dynamics within the MR1 binding pocket. Using root mean square fluctuation (RMSF) of each atom within the ligand, we quantified this dynamic motion (Figure 8b-d, left) and found that in contrast to DMRL, the RMSF of atoms composing the deazalumazine core of DZ are greater; conversely, the RMSF of atoms composing the ribityl moiety of DZ are smaller than those of DMRL. When comparing DZ and 2’-deoxy-DZ, ablation of the 2’-hydroxy enhances plasticity of the ribityl moiety without much change to the dynamics of the deazalumazine core. Through alignment of the dynamics of each ligand by its cyclic core, we visualized the extent of these fluctuations in the ribityl chains (Figure 8b-d, right) for a representative simulation of each ligand (simulations DMRL2, DZ1, and 2’-deoxy-DZ1, i.e. ddZ1). Taking snapshots every 10 nanoseconds of simulated time, we observed that the ribityl moiety of DMRL and 2’-deoxy-DZ were quite dynamic, though that of DMRL sampled space mostly through transposition of the entire ribityl moiety, while that of 2’-deoxy-DZ sampled space by rotation of the 5’ hydroxyl (reflected in the RMSF patterns). The ribityl chain of DZ, on the other hand, was strikingly rigid in all three of the simulation replicates. Our observation of enhanced fluctuations of the core-distal ribityl moiety atoms due to 2’-deoxy modification (in comparison with DZ) is consistent with those seen in previous crystallographic studies with of ternary complexes with synthetic deoxyribityl-versions of 5-OP-RU, as well as with molecular dynamics simulations performed on those ternary structures [9]. Together, these data suggest that minor modifications to MR1 ligands can alter their dynamics in the MR1 pocket, which may affect TCR recognition and explain the TCR selectivity that we have observed.

**Figure 8.**
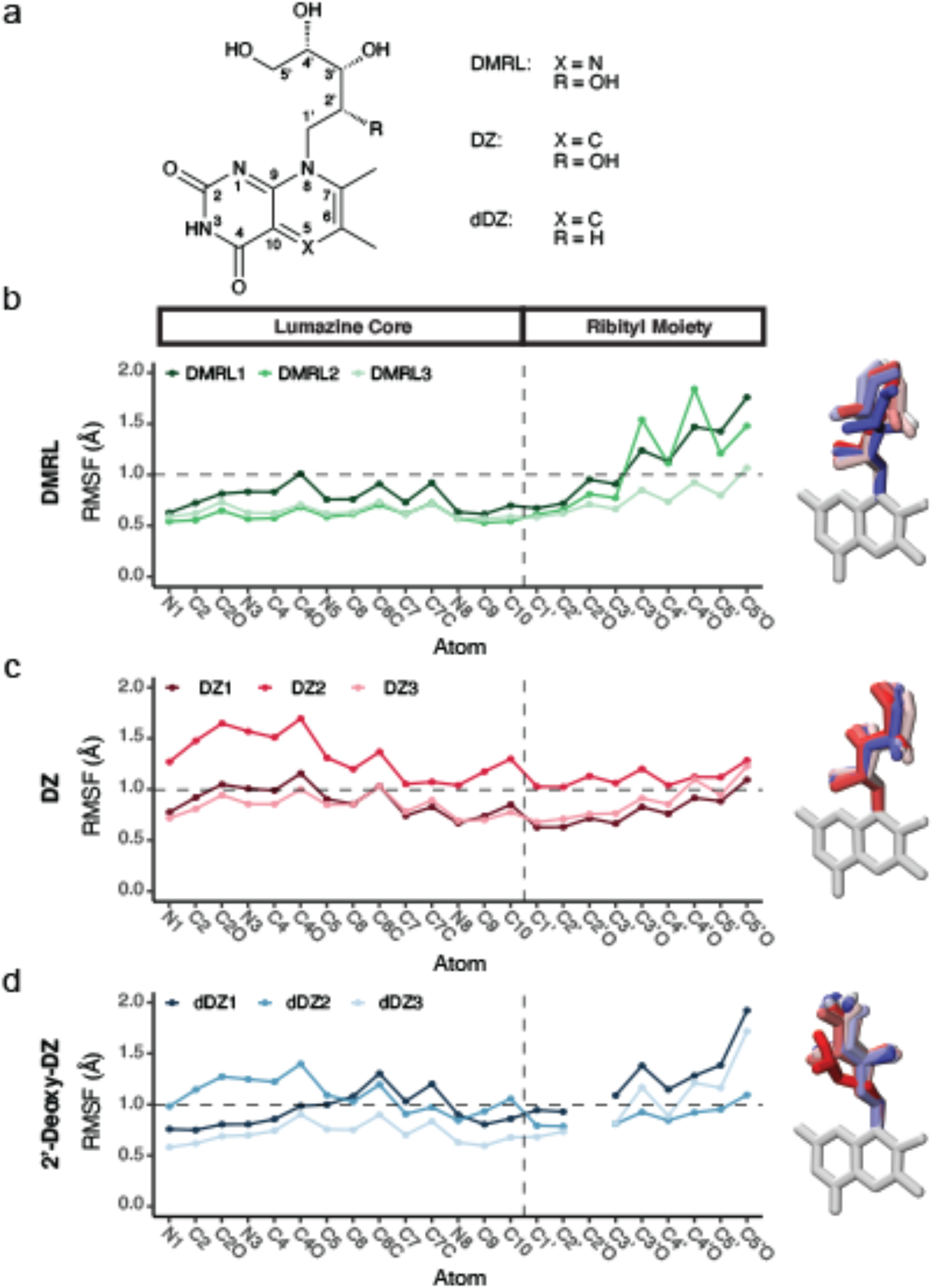
Molecular dynamics simulations of DZ, 2’-deoxy-DZ, and DMRL ligands docked to MR1 identify differential flexibility of each ligand within the binding pocket. a) Chemical representation of DMRL and its derivatives. b-d) Left panels show RMSF calculations (indicative of ligand dynamics) for each simulation replicate over the course of each 80 ns MD simulation for DMRL (b), DZ (c) or 2’-deoxy-DZ (dDZ) (d). The horizontal line at 1Å does not represent any parameter; it only serves as a visual guide to compare across plots. Right panels show visualization of dynamic states adopted by the ribityl chain in a representative simulation for each ligand (simulations DMRL2, DZ1, and dDZ1), evolving from red to blue across time.

## Discussion

Microbial metabolites presented on MR1 and recognized by MR1T cells represent a novel and largely unexplored class of T cell-activating ligands, but targeting MR1T cells for therapeutic or vaccine-related applications requires a better understanding of structure and stability of ligands interacting with MR1 and the MR1T TCR to induce activation. While pyrimidines like 5-OE-RU and 5-OP-RU are potent antigens, their instability *in vitro* makes them challenging candidates for these applications. Furthermore, while there are now multiple recent studies demonstrating that treatment with 5-OP-RU expands MR1T cells and protects against infection with organisms like *Francisella tularensis* and *Legionella longbeachii* in a mouse model (e.g. [29-31]), it is unclear whether these findings will translate to all organisms, or in a clinical setting. For example, in a mouse model, treatment with 5-OP-RU does not result in protection against *Mycobacterium tuberculosis*, despite similar MR1T cell expansion [32-34]. Additionally, a recent study in macaques demonstrated that treatment with 5-OP-RU not only failed to provide a therapeutic benefit in the context of *Mycobacterium tuberculosis*, it also resulted in functional impairment and exhaustion of MR1T cells. Together, these studies suggest that there is still much to understand with regard to the use of 5-OP-RU or other ligands for clinical purposes. The majority of the functional and molecular work done in the MR1 field has focused on 5-OP-RU and high affinity MAIT TCRs, leaving the ribityllumazine and other unidentified classes of ligands underexplored. For example, 26 of the 49 existing MR1:ligand-TCR structures contain 5-OP-RU or molecular variants of 5-OP-RU, and another 10 contain 6-FP or acetyl-6-FP, which are strong antagonists of MR1T cells. While useful tools for contrasting with other MR1T cell ligands, there is question as to the physiological relevance of these studies with 6-FP (and, some would argue, 5-OP-RU). In contrast, only three complex crystal structures are in complex with ribityllumazines. Further, 29 of the 49 crystal structures available are of the same TCR (known as “A-F7” or “F7”), leaving the diversity of MR1T cell TCR molecular recognition strategies largely unexplored. Thus, generation of synthetically modified ligands provides useful tools for both targeting of MR1T cells and for studying the complexity of the MR1:ligand-TCR interactions. Here, we reveal that even minor chemical changes to TCR-inaccessible positions of MR1 ligands affect antigenicity of these ligands, highlighting the high level of antigen selectivity exhibited by MR1-restricted T cells. Interestingly, certain MR1-restricted T cell clones were agnostic to these chemical modifications, indicating the importance of studying the MR1:ligand-TCR interaction closely.

Our data strengthen a model where minor ligand modifications affect antigen potency [9, 17, 23]. Previous work has shown that the potency of a deaza-5-OP-RU analogue was decreased considerably when assayed in an *in vitro* MR1T cell reporter assay with a Jurkat cell expressing a single MR1T TCR [17]. However, analysis of this molecule in the context of multiple MR1T cell TCRs could provide insight into whether the deaza-modification has different functional impacts for TCRs. When we made a similar modification to the 5-nitrogen of PLI to generate DZPLI, we also saw significantly decreased potency in the ability to activate MR1T cells, though this appears to be driven by failure to surface stabilize MR1 and not caused by TCR selectivity. The same was not true when we modified DMRL in the same way to generate DZ. While two of our MR1T TCRs exhibited a similar decrease in recognition of DZ, the deaza-version of DMRL was actually equally or more potent for three different MR1T TCRs. This was unexpected based on our results with DZPLI and those of Mak *et al* [17].

Since all of the TCRs described in this paper are capable of recognizing the highly potent 5-OP-RU ligand, these data clearly indicate that MR1T cells can discriminate between structurally similar ligands, though we do not yet understand the molecular determinants of this phenomenon. This is particularly striking because the derivatization of DMRL to DZ involves modification of the ligand at a location that is likely bound deep in the A’ pocket of MR1, and therefore inaccessible to the TCR. We find that one possible explanation of this observation is that modifications to the cyclic core may affect the dynamics of both MR1 and more distal atoms of the ribityl group, leading to differential CDR3 interactions with this group and therefore variability in MR1T cell reactivity. This is consistent with previous reports of simulations done on the MR1-TCR ternary complex with 2’-deoxy-5-OP-RU and 5’-deoxy-5-OP-RU, wherein the distal atoms of the ribityl moiety exhibited greater fluctuation in the simulations with the former [9]. However, there was no comparison with the parental 5-OP-RU ligand and simulation replicates were not performed in this study, making it hard to determine whether observation was due to stochastic effects. Together, these data provide support for the continued study of MR1 ligand modifications and the molecular mechanism of MR1T cell ligand discrimination.

Beyond unanswered questions with regard to the importance of observed differences in MR1T TCR recognition of novel and synthetically modified ligands *in vivo* are questions related to how these modifications may impact the ability of ligands to be adjuvanted or transported *in vivo*. Legoux *et al*. demonstrated that 5-OP-RU could rapidly reach the thymus of mice after being painted on the skin of the thorax [35], but the mechanism by which this happens is not yet known.

Delivery of 5-OP-RU in this way resulted in an increase in the numbers and activation of early MR1T cell precursors from the thymus [35]. However, data demonstrating the selective clonal expansion of MR1T cells with distinct TCRs following infection with different microbes [12], and vaccine studies with 5-OP-RU [29-34] suggest that there are likely differences from what happens naturally in the context of natural infection with diverse microbes producing discrete ligands. Furthermore, there may be differences in the types of ligands that are generated in the context of intracellular infection with microbes compared to exogenous delivery of synthetic ligands. Thus, it is not clear whether exogenous delivery of a ligand with broad and potent MR1T cell activation (e.g. 5-OP-RU) will be the optimal way to target MR1T cells for therapeutic or vaccine purposes. Taken together, our data demonstrate that analysis of multiple MR1T cell antigen analogues in the context of diverse MR1T T cell receptors will be important to improving our understanding of the stability and relationship to activation of MR1T cells.

### Experimental procedures

#### General Chemical synthesis

Reagents were obtained from Sigma-Aldrich or Fisher Scientific. ^1^H-NMR was recorded on a Bruker DPX spectrometer at 400 MHz. Chemical shifts are reported as parts per million (ppm) downfield from an internal tetramethylsilane standard or solvent references. High-resolution mass spectra were acquired on a ThermoElectron LTQ-Orbitrap Discovery high resolution mass spectrometer with a dedicated Accela HPLC system by Andrea DeBarber at the Bioanalytical MS facility, Portland State University. For air- and water-sensitive reactions, glassware was oven-dried prior to use and reactions were performed under argon. Dichloromethane, N,N-dimethylformamide, and tetrahydrofuran were dried using a solvent purification system manufactured by Glass Contour, Inc. (Laguna Beach, CA). All other solvents were of ACS chemical grade (Fisher Scientific) and used without further purification unless otherwise indicated. Analytical thin-layer chromatography was performed with silica gel 60 F_254_ glass plates (SiliCycle). Flash column chromatography was conducted with pre-packed normal or reversed phase columns (Biotage). High performance liquid chromatography (HPLC) was performed on an Agilent 1260 Infinity system with a flow rate of 1.0 mL/min using a Sunfire C18-A column 150 × 4.6 mm, 5 micron analytical column or a Sunfire 30 × 50 mm, 5 micron preparative column. HPLC analytical conditions: mobile phase (MP) A: 0.1% formic acid in water, B: 0.1% formic acid in acetonitrile (ACN), flow rate = 1.0 mL/min, gradient: 0% B for 2 min, 0-100% B over 13 min, 100% B for 2 min, UV-Vis detection at *λ*_1_ = 254 nm and *λ*_2_ = 220 nm. All final products were ≥95% purity as assessed by this method. Retention time (*t*_R_) and purity refer to UV detection at 220 nm. Preparative HPLC conditions: mobile phase (MP) A: 0.1% formic acid in water, B: 0.1% formic acid in acetonitrile (ACN), flow rate = 10.0 mL/min, gradient **A**: 0-30% B over 7 min, 30-50% B over 2 min, 100% B for 1 min; gradient **B**: 30% B for 6 min, 30-50% B over 8 min, 100% B for 4 min, UV-Vis detection at *λ*_1_ = 254 nm and *λ*_2_ = 220 nm.

#### 6-(((2S, 3S, 4R)-2,3,4,5-Tetrahydroxypentyl)amino)pyrimidine-2,4-(1H,3H)-dione (2a)

Commercially available ribitylamine [36] (150 mg, 1.0 mmol) and chlorouracil (42 mg, 0.28 mmol) were dissolved in 15 mL water in a 20 mL microwave vial and microwaved at 150°C for 2 h. After the reaction was complete, the desired ribitylaminouracil product (24 mg, 0.092 mmol, 33% yield) was isolated by preparative HPLC using gradient A. ^1^H-NMR was consistent with reference spectra [2].

#### 6-(((3S,4R)-3,4,5-Trihydroxypentyl)amino)pyrimidine-2,4-(1H,3H)-dione (2b)

Commercially available 1-amino-1,2-dideoxy D-*erythro*-pentitol [37] (135 mg, 1 mmol) and chlorouracil (42 mg, 0.28 mmol) were dissolved in 15 mL water in a 20 mL microwave vial and microwaved at 150°C for 2 h. After the reaction was complete, the desired aminouracil product (26 mg, 0.11 mmol, 39% yield) was isolated by preparative HPLC using gradient A and used without further purification.

#### 6,7-Dimethyl-8-D-ribityldeazalumazine (DZ)

Ribitylaminouracil **2a** (26 mg, 0.10 mmol) and the sodium salt of commercially available 2-methyl butan-3-one-ol [38] (26 mg, 0.20 mmol) were refluxed in 0.5 M HCl (1.0 mL) at 100°C for 2 h. After the reaction was complete, DZ (8.0 mg, 0.025 mmol, 25 % yield) was isolated by preparative HPLC using gradient B. ^1^H-NMR was consistent with reference spectra [20]. ^1^H-NMR (400 MHz, D_2_O) δ 2.35 (s, 3H), 2.74 (s, 3H), 3.64 (m, 2H), 3.68 (m, 2H), 3.84 (m, 2H), 4.31 (m, 2H), 4.92 (m, 2H), 8.40(s, 1H); HRMS (ESI-TOF) Calculated for C_14_H_20_N_3_O_6_ [M +H^+^] 326.1352, Found 326.1347.

#### 6,7-Dimethyl-8-D-(2’-deoxyribityl)deazalumazine (2’-deoxy-DZ)

Compound **2b** (26 mg, 0.10 mmol) and the commercially available sodium salt of 2-methyl butan-3-one-ol (26 mg, 0.20 mmol) were refluxed in 0.5 M HCl (1.0 mL) at 100°C for 2 h. After the reaction was complete, 2’-deoxy-DZ (8.0 mg, 0.025 mmol, 25 % yield) was isolated by preparative HPLC using gradient B. ^1^H-NMR (400 MHz, D_2_O/CD_3_OD) δ 1.93 (m, 2H), 2.19 (m, 2H), 2.41 (s, 3H), 2.74 (s, 3H), 3.34-3.74 (m, 5H), 8.31(s, 1H); HRMS (ESI-TOF) Calculated for C_14_H_20_N_3_O_5_ [M +H^+^] 310.1397, Found 310.1347.

#### 3-(2,4,7-Trioxo-8-((2S,3S,4R)-2,3,4,5-tetrahydroxypentyl)-1,2,3,4,7,8-hexahydropyrido[2,3-d]pyrimidin-6-yl)propanoic acid (DZPLI)

Ribitylaminouracil **2a** (26 mg, 0.1 mmol) and commercially available 1,5-diethyl-2-formylpentanedioate (75 mg, 0.40 mmol) were refluxed in 0.5 M HCl (1.0 mL) at 100°C for 2 h. The crude mixture was then hydrolyzed in 2 M LiOH at 40°C overnight. After the reaction was complete, DZPLI (4.2 mg, 0.011 mmol, 11% yield) was isolated by preparative HPLC using gradient B. ^1^H-NMR (400 MHz, CD_3_OD) δ 2.61 (t, 2H), 2.82 (t, 2H), 3.67 (m, 2H), 3.83 (m, 2H), 4.14 (m, 1H), 4.35 (m,1H), 4.64 (d, 1H), 7.84(s, 1H); HRMS (ESI-TOF) Calculated for C_15_H_20_O_9_N_3_ [M + H^+^] 386.1205, Found 386.1194.

#### 4-Chloro-2, 6-dimethoxypyrimidine-5-carbaldehyde (4)

An oven-dried flask was charged with commercially available 4-chloro-2,6-dimethoxypyrimidine (0.95 g, 5.5 mmol) and then evacuated and backfilled with argon. Anhydrous THF (5 mL) was added through a rubber septum. The mixture was cooled to -78°C and a 1.6 M solution of n-butyllithium (n-BuLi) in hexanes (3.8 mL, 6.0 mmol) was added dropwise. The mixture was stirred for an additional 0.5 h, and DMF (1 mL, 13 mmol) was added and stirring was continued for 2 h at the same temperature. The reaction was quenched by addition of aqueous HCl (1.6 M, 25 mL), and the mixture was extracted with ether (3 × 40 mL). The combined organic layers were washed with aqueous HCl (1.6 M, 25 mL) and water (40 mL), dried (Na_2_SO_4_) and evaporated to dryness. The residue was purified by column chromatography on silica gel (ethyl acetate-toluene, 1:6) to afford 4-chloro-2,6-dimethoxypyrimidine-5-carbaldehyde **4** (0.80 g, 4.0 mmol, 73%). ^1^H-NMR was consistent with reference spectra [39]. ^1^H-NMR (400 MHz, CDCl_3_) δ 4.11 (s, 3H), 4.15 (s, 3H), 10.34 (s, 1H).

#### (E)-4-(4-Chloro-2,6-dimethoxypyrimidin-5-yl)but-3-en-2-one (5)

To a solution of **4** (120 mg, 0.60 mmol) in toluene (4 mL) was added commercially available 1-(triphenylphosphoranylidene)-2-propanone (189 mg, 0.60 mmol). The reaction mixture was refluxed for 6 hours. After the reaction was complete, the organic solvent was removed *in vacuo*. The resulting residue was purified by column chromatography on silica gel (ethyl acetate-hexane, 1:3) to afford the desired product **5**. ^1^H-NMR (400 MHz, CDCl_3_) δ2.37 (s, 3H), 4.01 (s, 3H), 4.09 (s, 3H), 7.03 (d, 1H, *J*=18Hz), 7.67 (d, 1H, *J=*18Hz).

#### (E)-4-(2,4-Dimethoxy-6-(((2S,3S,4R)-2,3,4,5-tetrahydroxypentyl)amino)pyrimidin-5-yl)but-3-en-2-one (6)

To a stirred solution of **5** (63 mg, 0.26 mmol) in DMF (4 mL) was added ribitylamine (120 mg, 0.80 mmol). The reaction mixture was refluxed for 12 h. After the reaction was complete, the organic solvent was removed in vacuo. The resulting residue was purified by preparative HPLC to deliver the desired product **6** (21 mg, 0.060 mmol, 23% over two steps). ^1^H-NMR (400 MHz, CD_3_OD) δ2.33 (s, 3H), 3.95-3.51 (m, 9H), 4.01 (s, 3H), 4.18 (s, 3H), 6.83 (d, 1H, *J*=16.4), 7.48 (d, 1H, *J*= 16.4).

#### 7-Methyl-8-((2S,3S,4R)-2,3,4,5-tetrahydroxypentyl)pyrido[2,3-d]pyrimidine-2,4(3H,8H)-dione (monomethyl-DZ)

To a solution of **6** (20 mg, 0.050 mmol) in 6 M HCl (1 mL) was refluxed at 70°C overnight. After the reaction was complete, **monomethyl-DZ** (6.0 mg, 0.020 mmol, 39%) was purified by preparative HPLC. ^1^H-NMR was consistent with reference spectra [20]. ^1^H-NMR (400 MHz, D_2_O) δ2.78 (s, 3H), 3.41-3.88 (m, 5H), 4.29 (m, 1H), 4.44-4.65 (m, 5H) 7.18 (d, 1H, *J*=10.3), 7.48 (d, 1H, *J*= 12.2)

#### Human subjects

This study was conducted according to the principles expressed in the Declaration of Helsinki. Study participants, protocols and consent forms were approved by the Institutional Review Board at Oregon Health & Science University (OHSU) (IRB00000186).

#### Reagents and cells

Dendritic cells (DC) were derived from human peripheral blood monocytes as previously described [40, 41]. The bronchial epithelial cell line BEAS-2B (CRL-9609) was originally obtained from ATCC and was cultured in DMEM+10% heat inactivated fetal bovine serum (FBS). The BEAS-2B:ΔMR1 cell line was derived by CRISPR/Cas9 disruption of the MR1 gene, and MR1 expression was reconstituted in these cells [42]. Wild-type BEAS-2B cells overexpressing MR1 fused to GFP were previously described [22]. MAIT cell clones were derived, expanded, and maintained as previously described [11, 43].

#### ELISPOT assay

DC or BEAS-2B cells were harvested, counted and used in equivalent numbers, as indicated in the figure legends, as antigen presenting cells in an ELISPOT assay with IFN-γ production by MAIT cell clones as the readout as previously described [43]. Synthetic compounds or positive controls (*M. smegmatis* supernatant or PHA) were added to the cells at concentrations indicated for one hour prior to addition of the MAIT cell clones, and the ELISPOT plates were incubated for 18 hours prior to development. Phytohemagglutinin (PHA) was used at 10 μg/ml. Supernatant from *M. smegmatis* was prepared by collecting the <3 kDa fraction of supernatant from logarithmically-growing bacteria using a size exclusion column (Millipore). The volume of supernatant required for maximal response in the assay was determined empirically following preparation. Blocking was performed using the α‐MR1 26.5 clone (Biolegend) and an IgG2a isotype control, added at 2.5 μg/ml for 1 hour prior to the addition of ligand.

#### Flow cytometry

BEAS-2B:MR1-GFP cells were grown in a 6-well tissue culture plate to ∼70% confluency, and then incubated with synthetic compounds or vehicle at the indicated concentrations for 16 hours. Cells were harvested on ice and surface stained with the anti-MR1 26.5 antibody (1:100) conjugated to APC (Biolegend) for 40 minutes on ice in the presence of 2% human serum, 2% goat serum, and 0.5% FBS. Cells were washed and fixed, and subsequently analyzed with a BD FACS Symphony flow cytometer and FACS Diva software (BD). All analyses were performed using FlowJo software (TreeStar).

#### MR1 ligand docking

The crystal structures of MR1 chosen for this analysis are contained within Protein Data Bank entry 4GUP [2], the crystal structure of the heterodimer of human MR1 C262S and human β_2_m bound to the MR1 ligand 6-formylpterin (6-FP), and 4L4V, which contains the same protein species, except they are in complex with 6-methyl-7-hydroxyl-8-D-ribityllumazine (HMRL) and a MAIT TCR [8]. For PDB 4GUP, chains A (MR1) and B (β_2_m), which together compose one of the two conformations of MR1/β_2_m in the asymmetric unit, were selected due to this conformation’s similarity to that found in structures of MR1:ligand-TCR complexes (PDB ID: 4L4V, 4IIQ, 6PUF [5, 8, 9]). For PDB 4L4V, chains A (MR1) and B (β_2_m) were also chosen, though there was only minor structural heterogeneity between the two copies in the asymmetric unit. For 4LCC, a structure of chimeric human-bovine MR1 in complex with rRL-6-CH_2_OH and the same MAIT TCR as that of 4L4V, chain A (single chain β_2_m-MR1) was chosen. The PDBs were stripped of the remaining polypeptide chains, the ligands, all waters, and crystallographically-resolved ions. Separately, HMRL, DMRL, DZ, 2’-deoxy-DZ, monomethyl-DZ, rRL-6-CH_2_OH, PLI, and DZPLI ligands were sketched with ChemDraw 18.2 and copied to BIOVIA Discovery Studio [26] for exporting as Mol2 files with 3D information. Ligand files were subsequently converted to PDB format using Pymol [44] and visually inspected for appropriate geometry and bond angles. PDBQT files were prepared for both protein and ligand files using AutoDockTools suite [45, 46], which provides additional information regarding partial charge, atom type, and rotatable bonds. In order to restrict docking to the A’ pocket, Vina [25] was run using an *x, y, z* box size of 18, 20, 14 Å centered at *x, y, z* coordinates -6.19, -9.44, -11.53. All other Vina parameters were set to the default. Each ligand’s top binding mode was selected as the representative structure, and a single PDB was prepared for each dock in Pymol by combining MR1 and ligand output files. Alignments between the two donor structures were performed by aligning the C_α_ (alpha carbons) in the β sheet of the α_1_/α_2_ domains.

#### CHARMM input generation and molecular dynamics

The β_2_m/MR1:ligand complexes were solvated with the TIP3P water model using CHARMM-GUI Solution Builder [47-49], and neutralized with K^+^ and Cl^−^ ions at physiological concentrations (0.15 M). To obtain missing parameters, ligands were parametrized by PDB coordinates using CGenFF [50]. Generated inputs were uploaded to the Midway compute cluster of the University of Chicago Research Computing Center to execute MD simulations. Each simulation was allowed 0.5 ns of equilibration followed by 80 ns of production, and each ligand was simulated in triplicate for a total of 15 simulations. Equilibrated systems use an NVT ensemble and production runs use an NPT ensemble, with the temperature kept constant at 300.15° K using Langevin dynamics [51]. The simulations were kept at constant pressure at one bar with the Nosé–Hoover Langevin piston by allowing the cell box size to change semi-isotropically [52]. van der Waals interactions were computed using a Lennard-Jones force-switching function over 10–12 Å while long-range electrostatics used particle mesh Ewald [53]. Production runs used a 2-fs time step and the SHAKE algorithm to constrain the bonds having hydrogen atoms [54].

#### TCR contact modelling

TCRs were selected with help from the TCR3D database [55], sampling from a diverse repertoire of TRBV genes, including 6-1, 6-2, 6-4, and 20-1 (PDB ID: 4L4V, 4IIQ, 4L9L, 4PJ7, 4PJ8 [5-8]). The 25^th^ frame of the DZ and 2’-deoxy-DZ simulations were isolated and aligned to the 4GUP donor structure by the C_α_ (alpha carbons) in the β sheet of the α_1_/α_2_ domains of the heavy chain. Each TCR was then independently coordinated to the interface in the same fashion and CDR loops identified by IMGT V-Quest [56] were subsequently isolated for visualization.

#### Simulation analysis

Raw simulation data was processed using Bio3D [57], an R library with the ability to read, write and process biomolecular structure and trajectory data. Root mean square fluctuation (RMSF) was calculated using included functions to determine the conformational variance of each atom with respect to their mean position. Structural visualizations and alignments were performed using VMD [58], and renders were generated with Tachyon internal-memory processes. Ribityl time lapse was accomplished by aligning dynamics by aromatic core, and displaying the initial ring structure while selecting the ribityl pose every 8 ns starting at 3 ns (to allow for equilibration). RMSD and RMSF were plotted using the ggplot2 library in R [59].

#### Data analysis

Unless otherwise indicated, experimental data were plotted and analyzed for statistical significance using Prism 9 (GraphPad Software).

## Data Availability

PDB coordinates for docked ligands have been uploaded as Supporting Information and are freely available. Description of the contents of each file can be found in the Supporting Information document. All other data are contained within the manuscript.

## Author contributions

All authors contributed to conception and design of studies, analysis and interpretation of data; HJ, AMP, NAL, MC, MN contributed to acquisition of data; HJ, AN, NAL and MJH wrote the manuscript. All authors contributed to revisions and approved the final version of the manuscript.

