## Supporting Information for "Deaza-modification of MR1 ligands modulates recognition by MR1-restricted T cells"

**Contents:**

**Figure S1:** Comparisons of docking modes in MR1 donor structures

**Figure S2:** Comparison of docked ligands to crystallographically-determined ligand binding mode

**Figure S3:** RMSD of ligands during simulation trajectories

**PDB Files:** Coordinates for donor structures and docked ligands

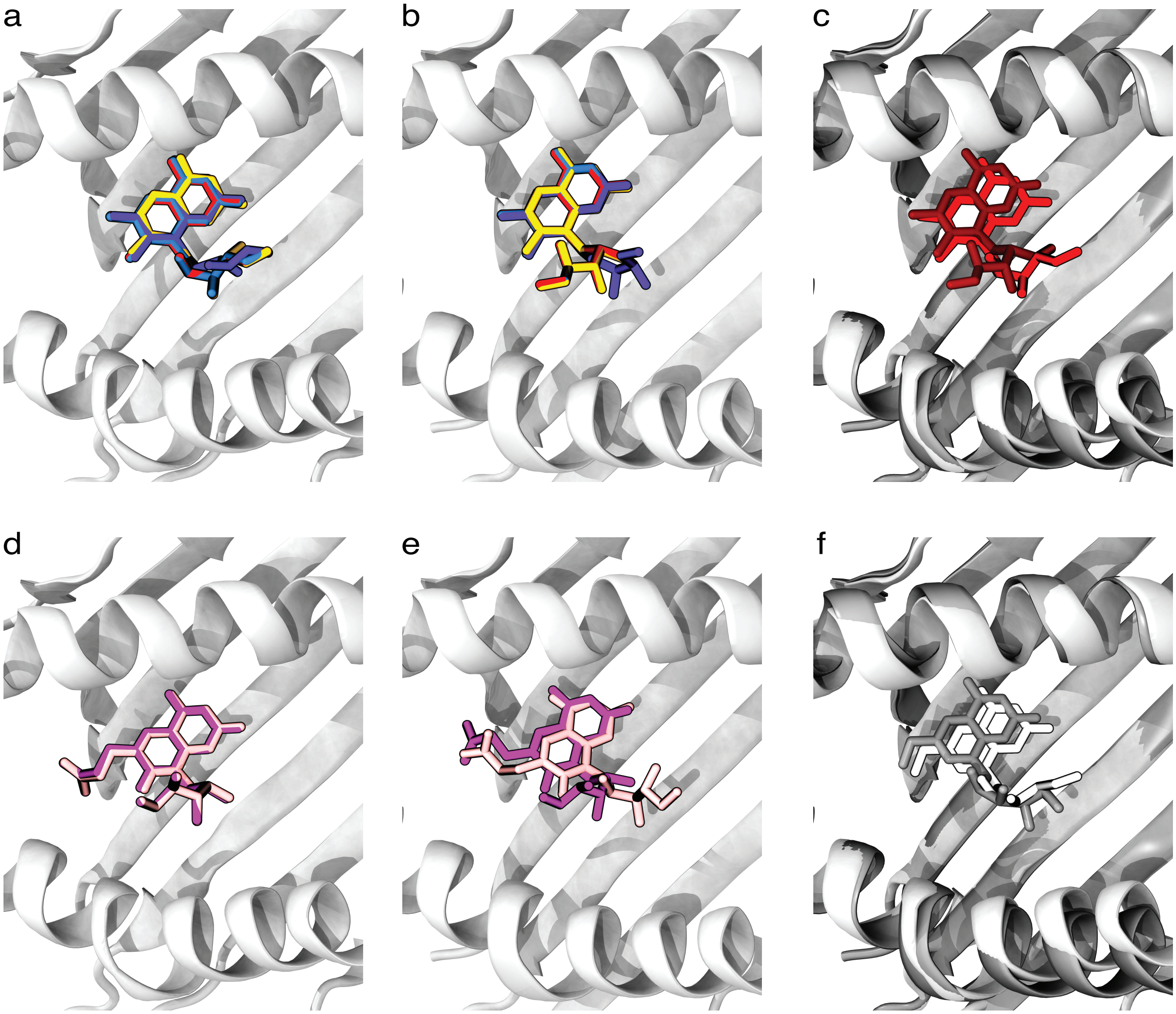


**Figure S1.** Docking comparisons for relevant donor structure/ligand pairs. a-b) DMRL (red), DZ (blue), 2'-deoxy-DZ (purple) and monomethyl-DZ (yellow) docked into 4GUP (a) and 4L4V (b). c) Comparison of the docking mode for DMRL in 4GUP (red) and 4L4V (maroon). d-e) PLI (mauve) and DZPLI (light pink) docked into 4GUP (d) and 4L4V (e). f) Comparison of the docking mode for rRL-6-$CH_2OH$ in 4GUP (white) and 4L4V (gray).

a

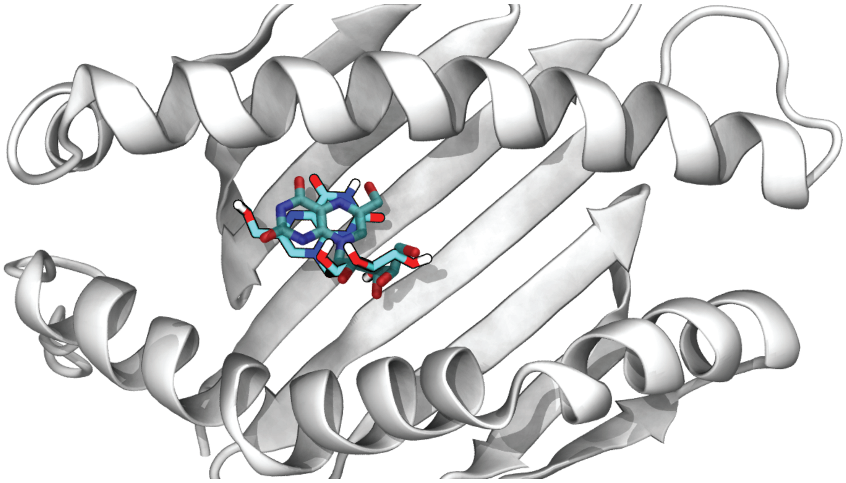

b

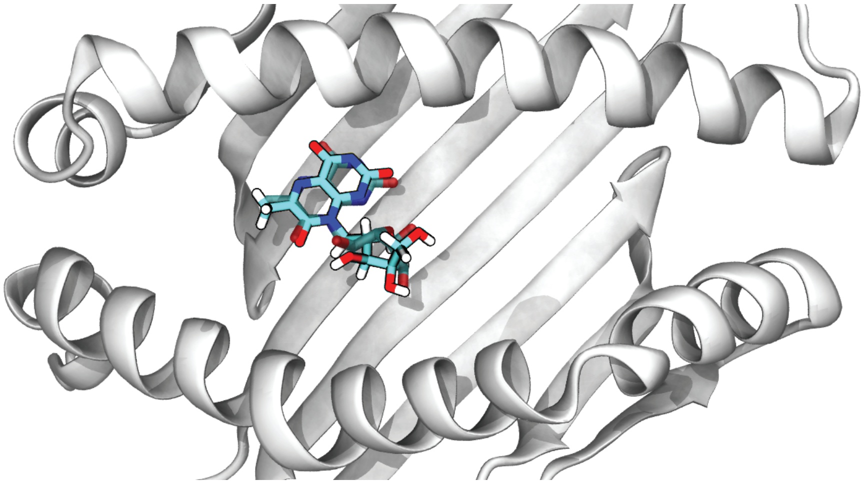

**Figure S2.** Comparisons between docking results and experimentally determined structures of ligand-MR1 complexes. The crystal structure ligand is shaded (dark cyan) and the docked ligand structure is vibrant (cyan).  a) rRL-6-$CH_2OH$ docked into 4LCC remains coplanar with the experimentally determined structure, but adopts a conformation ~180° flipped relative to that seen in the crystal structure. b) HMRL docked into 4L4V shows good agreement with that seen in the crystal structure.

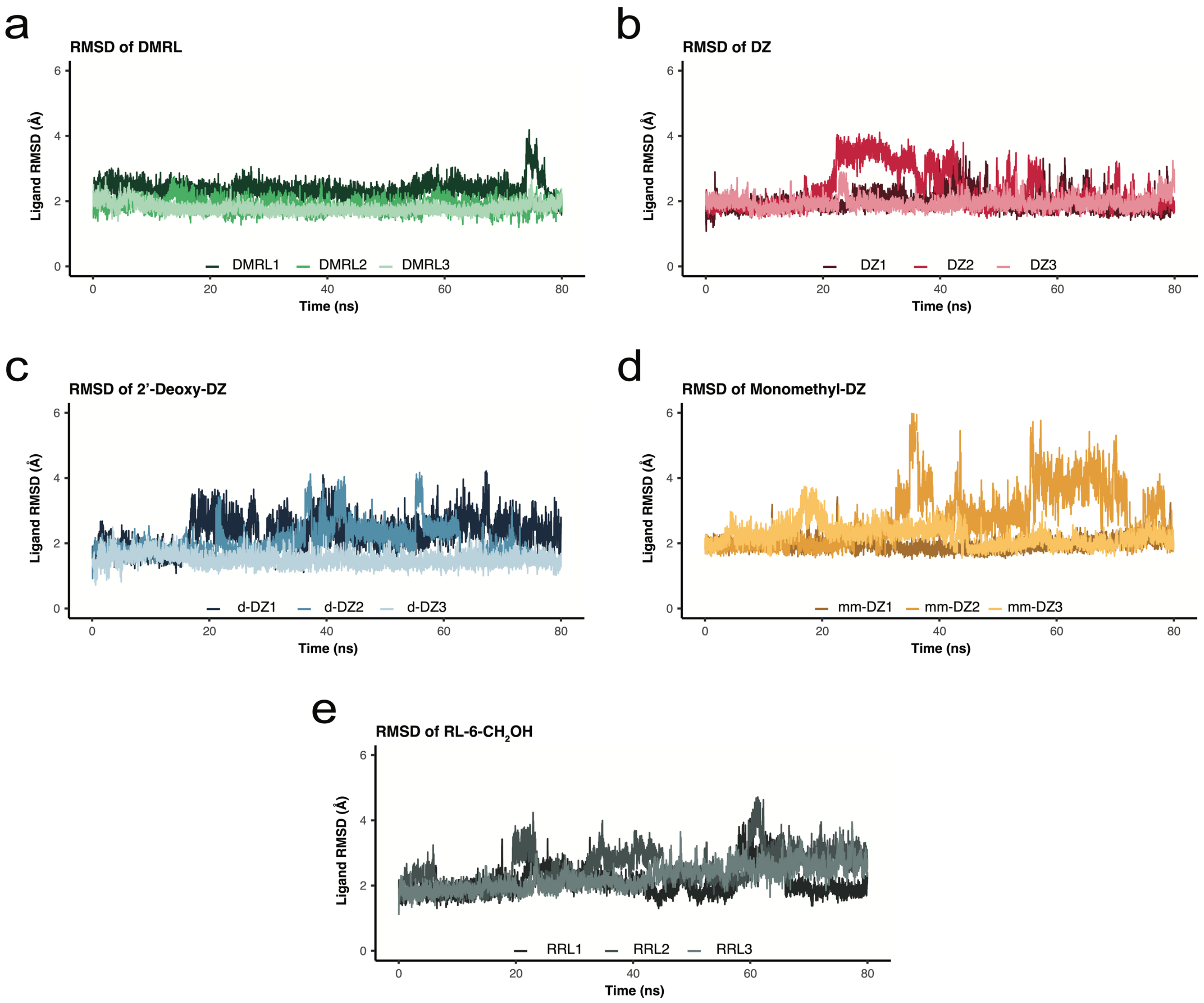


**Figure S3.** Root mean square deviation (RMSD) calculations of each ligand within the binding pocket of MR1 highlight their stability within the pocket over the course of each simulation. Full traces of this RMSD are shown for a) DMRL, b) DZ, c) 2'-deoxy-DZ, d) monomethyl-DZ, e) rRL-6-$CH_2OH$.

**PDB Files.** When files are opened in the same session in a software used for viewing molecular coordinates (e.g. Pymol or VMD), they will open as aligned for analyses in the main manuscript.

4gup.pdb: Donor structure of MR1 (originally crystallized with 6-FP)

4gup_dmrl.pdb: DMRL docked into the 4GUP donor structure

4gup_dz.pdb: DZ docked into the 4GUP donor structure

4gup_ddz.pdb: 2'-Deoxy-DZ docked into the 4GUP donor structure

4gup_mmdz.pdb: Monomethyl-DZ docked into the 4GUP donor structure

4gup_pli.pdb: PLI docked into the 4GUP donor structure

4gup_dzpli.pdb: DZPLI docked into the 4GUP donor structure

4gup_rrl.pdb: rRL-6-$CH_2OH$ docked into the 4GUP donor structure

4l4v.pdb: Donor structure of MR1 (originally crystallized with a ribityllumazine and a TCR)

4l4v_dmrl.pdb: DMRL docked into the 4L4V donor structure

4l4v_dz.pdb: DZ docked into the 4L4V donor structure

4l4v_ddz.pdb: 2'-Deoxy-DZ docked into the 4L4V donor structure

4l4v_mmdz.pdb: Monomethyl-DZ docked into the 4L4V donor structure

4l4v_pli.pdb: PLI docked into the 4L4V donor structure

4l4v_dzpli.pdb: DZPLI docked into the 4L4V donor structure

4l4v_rrl.pdb: rRL-6-$CH_2OH$ docked into the 4L4V donor structure

4l4v_hmrl_docked.pdb: HMRL docked into the 4L4V donor structure

4l4v_hmrl_crystallographic.pdb: crystallographically-determined docking mode of HMRL
